# Inverse Problem Reveals Conditions for Characteristic Retinal Degeneration Patterns in Retinitis Pigmentosa under the Trophic Factor Hypothesis

**DOI:** 10.1101/2021.09.04.458968

**Authors:** Paul A. Roberts

## Abstract

Retinitis pigmentosa (RP) is the most common inherited retinal dystrophy with a prevalence of about 1 in 4000, affecting approximately 1.5 million people worldwide. Patients with RP experience progressive visual field loss as the retina degenerates, destroying light-sensitive photoreceptor cells (rods and cones), with rods affected earlier and more severely than cones. Spatio-temporal patterns of retinal degeneration in human RP have been well characterised; however, the mechanism(s) giving rise to these patterns have not been conclusively determined. One such mechanism, which has received a wealth of experimental support, is described by the trophic factor hypothesis. This hypothesis suggests that rods produce a trophic factor necessary for cone survival; the loss of rods depletes this factor, leading to cone degeneration. In this paper we formulate a partial differential equation mathematical model of RP in one spatial dimension, spanning the region between the retinal centre (fovea) and the retinal edge (ora serrata). Using this model we derive and solve an inverse problem, revealing for the first time experimentally testable conditions under which the trophic factor mechanism will qualitatively recapitulate the spatio-temporal patterns of retinal regeneration observed in human RP.

## 1 INTRODUCTION

The group of inherited retinal diseases known as retinitis pigmentosa (RP) causes the progressive loss of visual function (Hamel, 2006; Hartong et al., 2006). The patterns of visual field loss associated with the human version of this condition have been well characterised (Grover et al., 1998); however, the mechanisms underpinning these patterns have yet to be conclusively determined (Newton and Megaw, 2020). In this paper, we use mathematical models to predict the conditions under which a trophic factor mechanism could explain these patterns.

The retina is a tissue layer lining the back of the eye containing light-sensitive cells known as photo-receptors, which come in two varieties: rods and cones (Fig. 1A). Rods confer monochromatic vision under low-light (scotopic) conditions, while cones confer colour vision under well-lit (photopic) conditions (Oyster, 1999). In RP, rod function and health are typically affected earlier and more severely than those of cones, with cone loss following rod loss. Rods are lost since either they or the neighbouring retinal pigment epithelium express a mutant version of one or both alleles (depending on inheritance mode) of a gene associated with RP (over 80 genes have been identified to date, see Gene Vision and Birtel et al., 2018; Coussa et al., 2019; Ge et al., 2015; Haer-Wigman et al., 2017). It is hypothesised that cones are lost following rods since they depend upon rods either directly or indirectly for their survival (Daiger et al., 2007; Hamel, 2006; Hartong et al., 2006).

**Figure 1.**
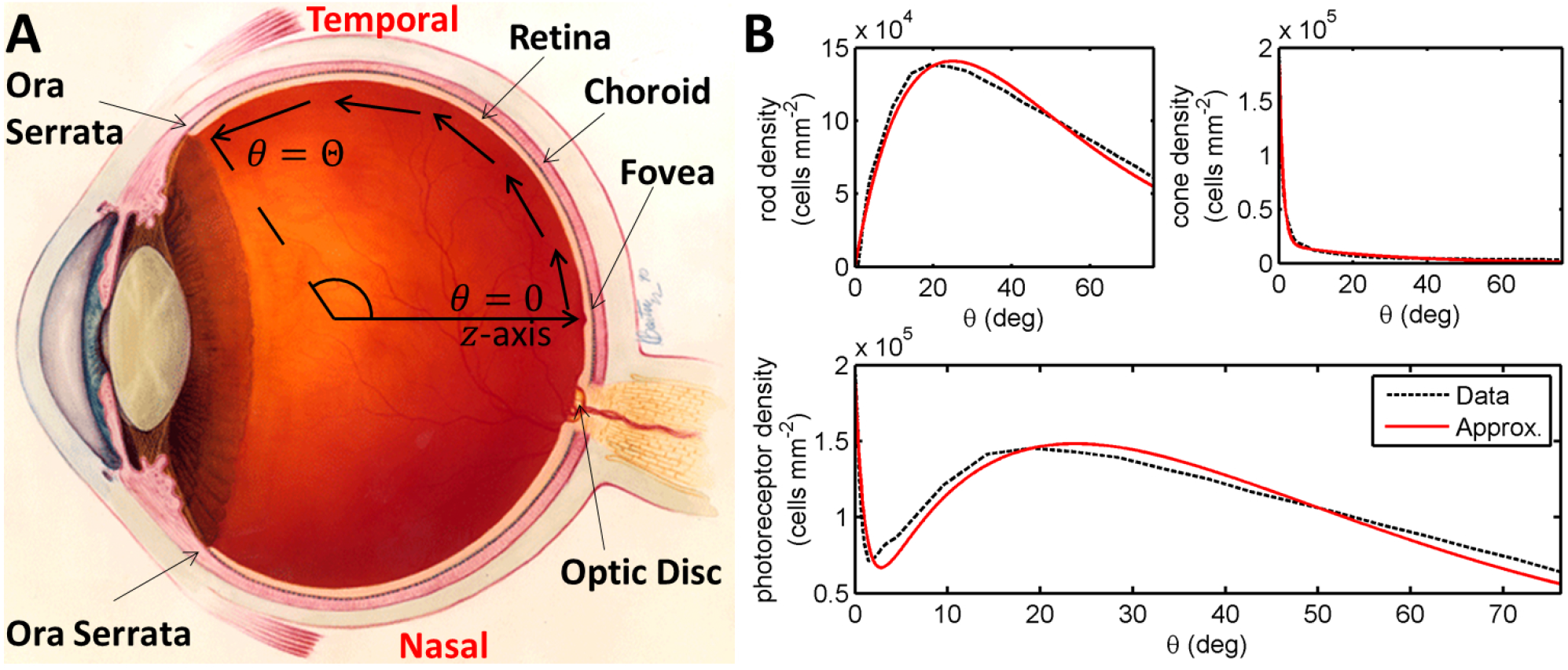
Diagrams of the human eye and retinal photoreceptor distribution (reproduced, with permission, from Roberts et al., 2017). **(A)** Diagram of the (right) human eye, viewed in the transverse plane, illustrating the mathematical model geometry. The model is posed on a domain spanning the region between the foveal centre, at *θ* = 0, and the ora serrata, at *θ* = Θ, along the temporal horizontal meridian, where *θ* measures the eccentricity. Figure originally reproduced, with modifications, from http://www.nei.nih.gov/health/coloboma/coloboma.asp, courtesy: National Eye Institute, National Institutes of Health (NEI/NIH). **(B)** Measured and fitted photoreceptor profiles, along the temporal horizontal meridian, in the human retina. Cone profile: 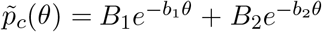, and rod profile: 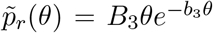 (see Table 2 for dimensionless parameter values). The photoreceptor profile is the sum of the rod and cone profiles 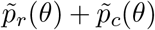. Experimental data provided by Curcio and published in Curcio et al. (1990).

A number of mechanisms have been hypothesised to explain secondary cone loss, including trophic factor (TF) depletion (Åt-Ali et al., 2015; Léveillard et al., 2004; Mei et al., 2016), oxygen toxicity (Stone et al., 1999; Travis et al., 1991; Valter et al., 1998), metabolic dysregulation (Punzo et al., 2009, 2012), toxic substances (Ripps, 2002) and microglia (Gupta et al., 2003). While not typically related to spatio-temporal patterns of retinal degeneration in the literature, it is reasonable to infer that these mechanisms play an important role in determining spatio-temporal patterns of retinal degeneration.

Grover et al. (1998) have classified the spatio-temporal patterns of visual field loss in RP patients into three patterns and six sub-patterns (see Fig. 2). Pattern 1A consists in a restriction of the peripheral visual field, while Pattern 1B also includes a para-/peri-foveal ring scotoma (blind spot); Pattern 2 (A, B and C) involves an initial loss of the superior visual field, winding nasally or temporally into the inferior visual field; lastly, Pattern 3 starts with loss of the mid-peripheral visual field, before spreading into the superior or inferior visual field and winding around the far-periphery. In all cases central vision is the best preserved, though it too is eventually lost (Hamel, 2006; Hartong et al., 2006). Patterns of visual field loss and photoreceptor degeneration (cell loss) are directly related (Escher et al., 2012), loss of the superior visual field corresponding to degeneration of photoreceptors in the inferior retina and vice versa, and loss of the temporal visual field corresponding to degeneration of photoreceptors in the nasal retina and vice versa.

**Figure 2.**
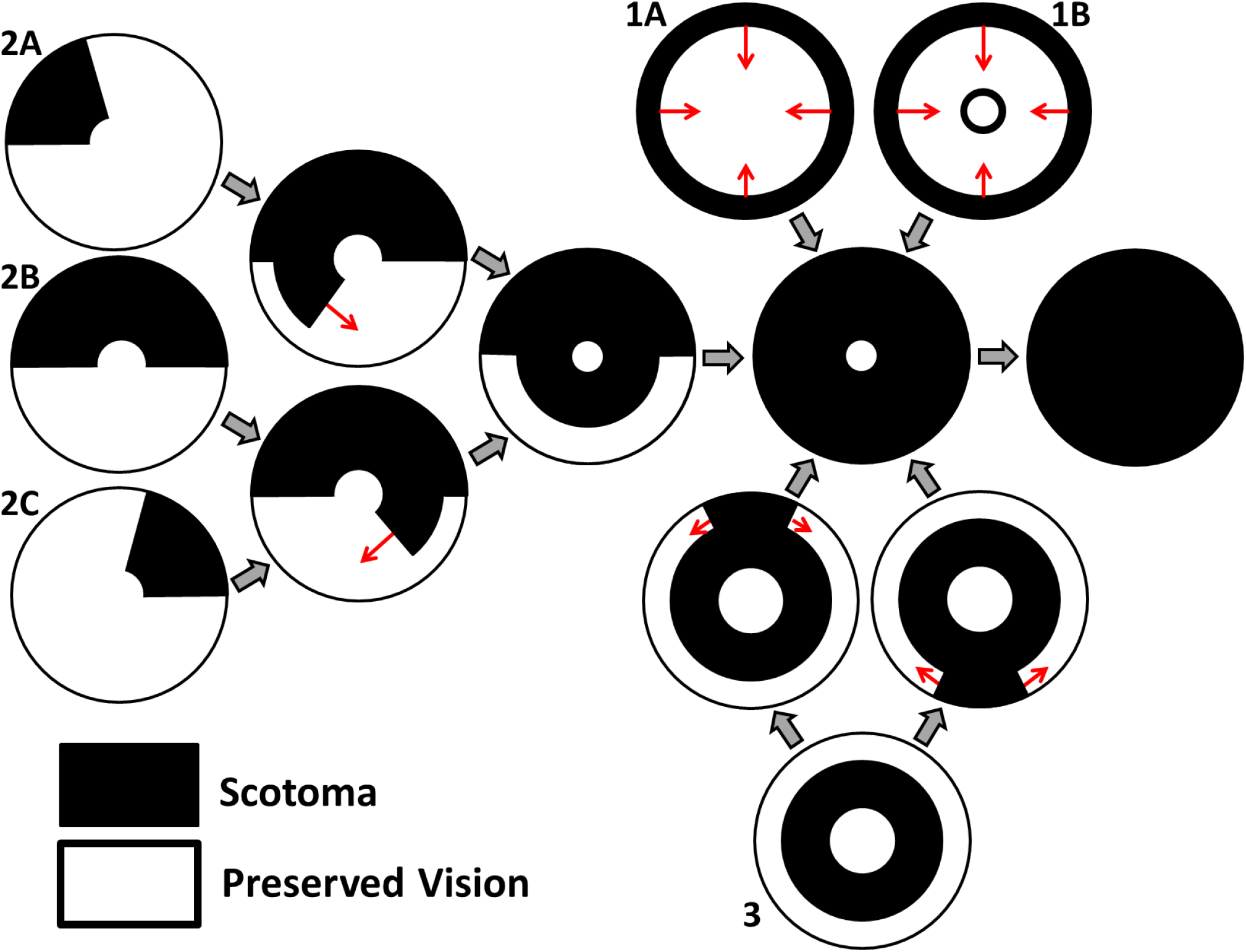
Characteristic patterns of visual field loss in human RP (reproduced, with permission, from Roberts et al., 2018). Visual field loss patterns can be classified into three cases and six subcases (classification system described in Grover et al., 1998). Large grey arrows indicate transitions between stages of visual field loss and small red arrows indicate the direction of scotoma (blind spot) propagation. See text for details.

In this paper we explore the conditions under which the TF mechanism, in isolation, can replicate the patterns of cone degeneration observed *in vivo*. Isolating a mechanism in this way enables us to identify the effects for which it is sufficient to account, avoiding confusion with other mechanistic causes. Understanding the mechanisms of secondary cone degeneration is important since it is the cones that provide high-acuity colour vision, and hence their loss, rather than the preceding rod loss, which is the most debilitating. Therefore, by elucidating these mechanisms, we can develop targeted therapies to prevent or delay cone loss, preserving visual function. The TF mechanism has been studied in detail. Rod photoreceptors have been shown to produce a TF called rod-derived cone viability factor (RdCVF), which is necessary for cone survival (Fintz et al., 2003; Léveillard et al., 2004; Mohand-Saïd et al., 1998, 2000, 1997; Yang et al., 2009). RdCVF increases cone glucose uptake, and hence aerobic glycolysis, by binding to the cone transmembrane protein Basigin-1, which consequently binds to the glucose transporter GLUT1 (Aït-Ali et al., 2015). Cones do not produce RdCVF, thus, when rods are lost, RdCVF concentration drops and cone degeneration follows (though it has been suggested that it may ultimately be oxygen toxicity which kills cones; Léveillard and Sahel, 2017).

Thus far, two groups have developed mathematical models operating under the TF hypothesis. Camacho *et al*. have developed a series of (non-spatial) dynamical systems ordinary differential equation models to describe the role of RdCVF in health and RP (Colón Vélez et al., 2003; Camacho et al., 2010; Camacho and Wirkus, 2013; Camacho et al., 2014, 2016a,b,c, 2019, 2020, 2021; Wifvat et al., 2021). In Roberts (2022), we developed the first partial differential equation (PDE) models of the TF mechanism in RP, predicting the spatial spread of retinal degeneration. It was found that, assuming all cones are equally susceptible to RdCVF deprivation and that rods degenerate exponentially with a fixed decay rate, the mechanism is unable to replicate *in vivo* patterns of retinal degeneration. Previous modelling studies have also considered the oxygen toxicity (Roberts et al., 2017, 2018 and related Roberts et al., 2016b) and toxic substance (Burns et al., 2002) mechanisms, predicting the spatio-temporal patterns of retinal degeneration they would generate. For a review of these and other mathematical models of the retina in health, development and disease see Roberts et al. (2016a).

In this study, we extend our work in Roberts (2022) by formulating and solving an inverse problem to determine the spatially heterogeneous cone susceptibility to RdCVF deprivation and rod exponential decay rate profiles that are required to qualitatively recapitulate observed patterns of spatio-temporal degeneration in human RP.

## 2 MATERIAL AND METHODS

### 2.1 Model Formulation

We begin by formulating a reaction-diffusion PDE mathematical model (a simplified version of the model presented in Roberts, 2022). Reaction-diffusion PDE models describe the way in which the spatial distribution of cells and chemicals change over time as a result of processes such as movement (diffusion), production, consumption, death and decay. We pose the model on a spherical geometry to replicate that of the human retina. This geometry is most naturally represented using a spherical polar coordinate system, (*r,θ,ϕ*), centred in the middle of the vitreous body, where *r ≥* 0 (m) is the distance from the origin, 0 *≤ θ ≤ π* (rad) is the polar angle and 0 *≤ ϕ <* 2*π* (rad) is the azimuthal angle. To create a more mathematically tractable model, we simplify the geometry by assuming symmetry about the *z*-axis (directed outward from the origin through the foveal centre), eliminating variation in the azimuthal direction, and effectively depth-average through the retina, assuming that it lies at a single fixed distance, *R >* 0 (m), from the origin at all eccentricities, *θ*, leveraging the fact that the retinal width is two orders of magnitude smaller than the eye’s radius (Oyster, 1999). Thus, we have reduced the coordinate system to (*R,θ*), where *R* is a positive constant parameter and 0 *≤ θ ≤* Θ is an independent variable, which we bound to range between the fovea (at *θ* = 0 rad) and the ora serrata (at *θ* = Θ = 1.33 rad; see Fig. 1A). We further simplify the model by non-dimensionalising; scaling the dependent and independent variables so that they and the resultant model parameters are dimensionless and hence unitless. This reduces the number of parameters (including eliminating *R*) and allows us to identify the dominant terms of the governing equations in the ensuing asymptotic analysis. For this reason, there are no units to be stated in Figs. 3–10. For the full dimensional model and non-dimensionalisation see Roberts (2022).

**Figure 3.**
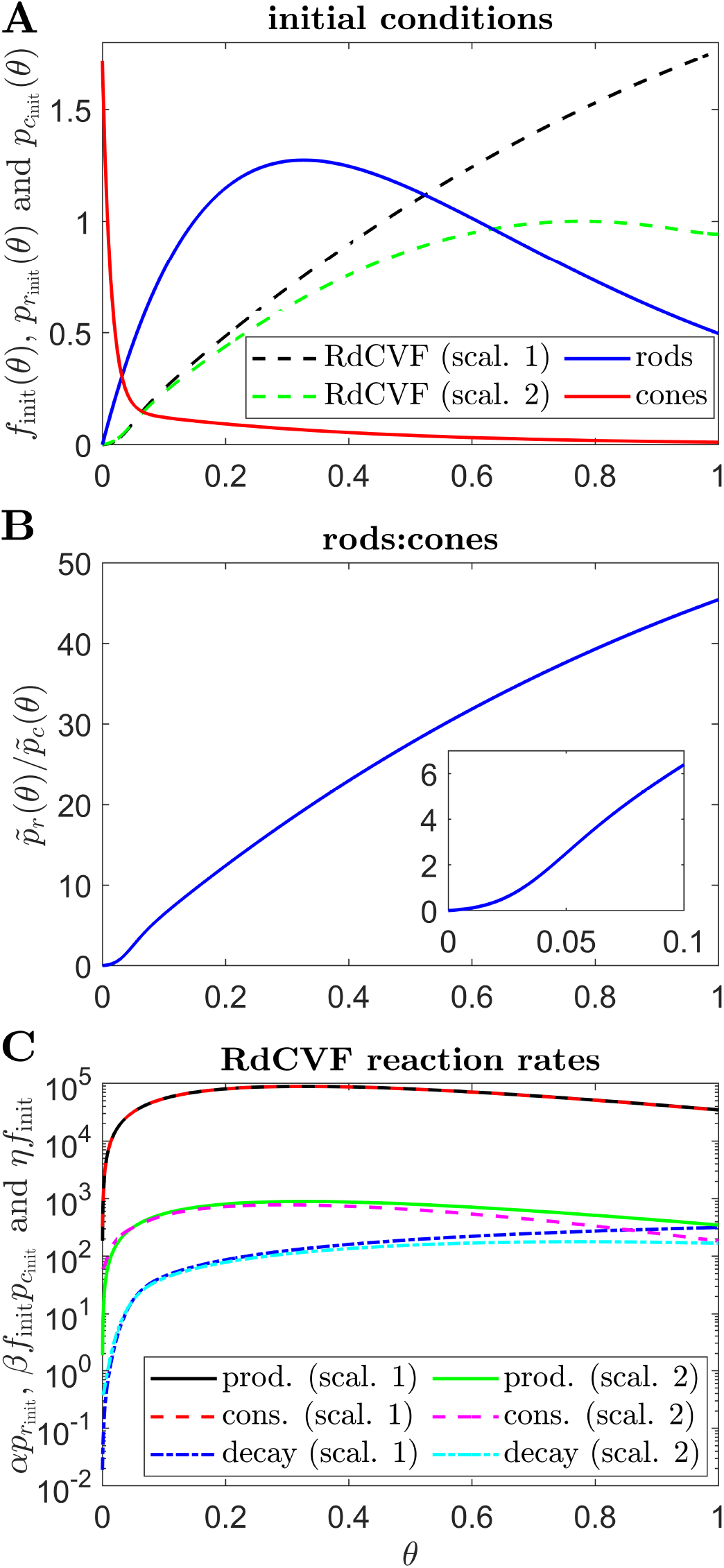
Initial conditions, ratio of rods to cones and RdCVF reaction rates. **(A)** initial conditions used in all simulations, consisting of healthy rod and cone profiles and the corresponding RdCVF profiles under Scalings 1 and 2 (the legend applies to **(A)** only). **(B)** variation in the healthy rod:cone ratio, 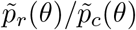, with eccentricity. **(C)** RdCVF production, consumption and decay rates under Scalings 1 and 2 (Eqn. (1), the legend applies to **(C)** only). To obtain *f*_init_(*θ*) in **(A)** and **(C)**, Eqs. (1) and (4) were solved at steady-state using the finite difference method, with 4001 mesh points, where 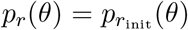 and 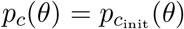. Under Scaling 1, *α* = 7.01 × 10^4^ and *β* = 1.79 × 10^6^, while under Scaling 2, *α* = 7.01 × 10^2^ and *β* = 1.79 × 10^4^. Remaining parameter values as in Table 2.

We proceed directly to the dimensionless model, which consists of a system of PDEs in terms of the dependent variables: TF concentration, *f* (*θ, t*), rod photoreceptor density, *p*_*r*_(*θ, t*), and cone photoreceptor density, *p*_*c*_(*θ, t*); as functions of the independent variables: polar angle, scaled to lie in the range 0 *≤ θ ≤* 1, and time, *t >* 0 (see Table 1).

**Table 1.** Variables employed in the non-dimensional mathematical model (Eqs. (1)–(5)).

| Variable | Description |
| --- | --- |
| $\theta$ | Eccentricity |
| $t$ | Time |
| $f(\theta, t)$ | Trophic factor concentration |
| $p_r(\theta, t)$ | Rod density |
| $p_c(\theta, t)$ | Cone density |

The TF equation is as follows

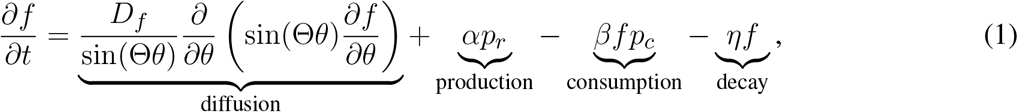

where *∂f/∂t* is the rate of change in TF concentration over time and the parameters, *D*_*f*_, the TF diffusivity, *α*, the rate of TF production by rods, *β*, the rate of TF consumption by cones, and *η*, the rate of TF decay, are positive constants. Trophic factor is free to diffuse across the retina through the interphotoreceptor matrix (Aït-Ali et al., 2015). We assume, in the absence of experimental evidence to the contrary, that all rods produce TF at an equal and constant rate, independent of the local TF concentration, such that the rate of TF production is directly proportional to the local rod density. Similarly, in the absence of further experimental evidence, we assume that all cones consume TF at an equal and constant rate for a given local TF concentration. Applying the physiological version of the Law of Mass Action, which states that the rate of a reaction is directly proportional to the product of the concentrations/densities of the reactants (Murray, 2002, in this case TF and cones), we assume that TF is consumed by cones at a rate directly proportional to the product of the local TF concentration and the local cone density. Lastly, we assume that TF decays exponentially, decreasing at a rate directly proportional to its local concentration, as has been shown to occur for a range of other proteins in living human cells (Eden et al., 2011).

The rod equation takes the following form

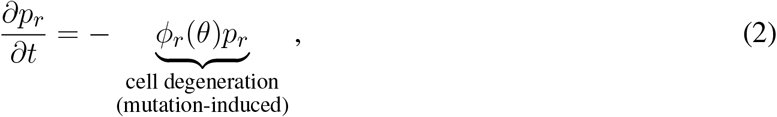

where *∂p*_*r*_*/∂t* is the rate of change in rod density over time and we allow the variable *ϕ*_*r*_(*θ*), the rate of mutation-induced rod degeneration, to vary spatially (functional forms defined in the Results section), or take a constant positive value, *ϕ*_*r*_. Rods degenerate due to their expression of a mutant gene (Hamel, 2006; Hartong et al., 2006) and are assumed to do so exponentially, at a rate directly proportional to their local density, consistent with measurements of photoreceptor degeneration kinetics in mouse, rat and canine models of RP (Clarke et al., 2000). Unlike with cones, RdCVF does not serve a protective function for rods (Aït-Ali et al., 2015); therefore, their rate of degeneration is independent of the TF concentration. We note that Eqn. (2) can be solved to yield 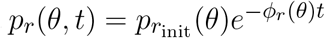 (where 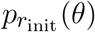, the initial value of *p*_*r*_(*θ, t*), is defined below), provided there is no delay in onset or interruption of degeneration.

The cone equation is as follows

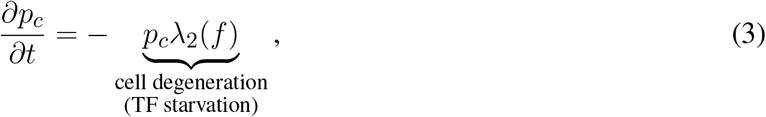

where *∂p*_*c*_*/∂t* is the rate of change in cone density over time. We define the Heaviside step function, *H*(·), such that

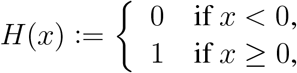

the function *λ*_2_(*f*) is given by

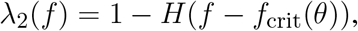

where we allow the variable *f*_crit_(*θ*), the TF threshold concentration, to vary spatially (functional forms defined in the Results section), or take a constant positive value, *f*_crit_. Cone density is assumed to remain constant provided the local TF concentration, *f* (*θ, t*), remains in the healthy range at or above the critical threshold, *f*_crit_, while cones are assumed to decay exponentially (due to TF starvation) at a rate directly proportional to their local density if *f* (*θ, t*) drops below this threshold, again consistent with Clarke et al. (2000)’s measurements of photoreceptor degeneration kinetics.

Having defined the governing equations (Eqs. (1)–(3)), we close the system by imposing boundary and initial conditions. We apply zero-flux boundary conditions at both ends of the domain,

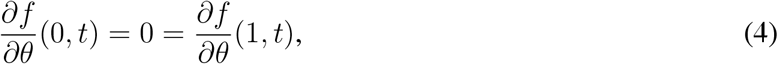

where *∂f/∂θ* is the TF concentration gradient in the polar direction, such that there is no net flow of TF into or out of the domain. This is justified by symmetry at *θ* = 0, while we assume that TF cannot escape from the retina where it terminates at the ora serrata (*θ* = 1). The healthy rod and cone distributions are given by the following functions

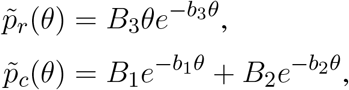

where the values of the positive constants *B*_1_, *B*_2_, *B*_3_, *b*_1_, *b*_2_ and *b*_3_ are found by fitting to the mean of Curcio et al. (1990)’s measurements of healthy human rod and cone distributions along the temporal horizontal meridian using the Trust-Region Reflective algorithm in Matlab’s curve fitting toolbox (see Fig. 1B). Lastly, we impose the initial conditions

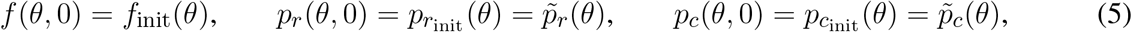

where *f*_init_(*θ*) is the steady-state solution to Eqs. (1) and (4) with 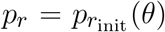 and 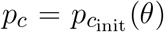 (see Fig. 3A). Thus, the retina starts in the healthy state in all simulations. See Table 2 for the dimensionless parameter values (see Roberts, 2022, for dimensional values and justification of parameter values). The model presented here simplifies that in Roberts (2022) in the following ways: it does not include treatment, cone outer segment regeneration, or initial patches of rod or cone loss, while mutation-induced rod loss is active for all simulations in this study. The present model also adds two new features to the previous model: allowing the rate of mutation-induced rod degeneration, *ϕ*_*r*_(*θ*), and the TF threshold concentration, *f*_crit_(*θ*), to vary spatially, where before they were constant (or piecewise constant in the high *f*_crit_ subcase).

**Table 2.** Parameters employed in the non-dimensional mathematical model (Eqs. (1)–(5)). Values are given to three significant figures (radians are dimensionless).

| Parameter | Description | Value |
| --- | --- | --- |
| $\Theta$ | Eccentricity of the ora serrata | 1.33 rad |
| $D_f$ | Trophic factor diffusivity | 0.237 |
| $\alpha$ | Rate of trophic factor production by rods | $7.01 \times 10^2$ or $7.01 \times 10^4$ |
| $\beta$ | Rate of trophic factor consumption by cones | $1.79 \times 10^4$ or $1.79 \times 10^6$ |
| $\eta$ | Rate of trophic factor decay | $1.79 \times 10^2$ |
| $\phi_r$ | Rate of mutation-induced rod degeneration | $7.33 \times 10^{-2}$ |
| $f_{\text{crit}}$ | Trophic factor threshold concentration | $3 \times 10^{-5}$ or 0.3 |
| $B_1$ | Cone profile parameter | 1.56 |
| $B_2$ | Cone profile parameter | 0.158 |
| $B_3$ | Rod profile parameter | 10.6 |
| $b_1$ | Cone profile parameter | 71.8 |
| $b_2$ | Cone profile parameter | 2.67 |
| $b_3$ | Rod profile parameter | 3.06 |

### 2.2 Numerical Solutions

Numerical (computational) solutions to Eqs. (1)–(5) were obtained using the method of lines (as in Roberts, 2022), discretising in space and then integrating in time. The time integration was performed using the Matlab routine ode15s, a variable-step, variable-order solver, designed to solve problems involving multiple timescales such as this (Matlab version R2020a was used here and throughout the paper). We used a relative error tolerance of 10^−6^ and an absolute error tolerance of 10^−10^, with the remaining settings at their default values. The number of spatial mesh points employed varies between simulations, taking values of 26, 51, 101, 401 or 4001. The upper bound of 4001 mesh points was chosen such that the distance between mesh points corresponds to the average width of a photoreceptor. In each case the maximum computationally feasible mesh density was employed, all mesh densities being sufficient to achieve accurate results. The initial TF profile, *f* (*θ*, 0) = *f*_init_(*θ*), was calculated by discretising Eqs. (1) and (4) at steady-state, using a finite difference scheme, and solving the consequent system of nonlinear algebraic equations using the Matlab routine fsolve (which employs a Trust–Region–Dogleg algorithm) with 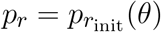 and 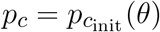.

### 2.3 Inverse Problem

Our previous modelling study of the TF hypothesis predicted patterns of cone degeneration which failed to match any known patterns in human RP (Roberts, 2022). In that study we made the simplifying assumption that model parameters are spatially uniform, such that they do not vary with retinal eccentricity. While this is a reasonable assumption in most cases, we have reason to believe that two of the parameters — the rate of mutation-induced rod loss, *ϕ*_*r*_, and the TF threshold concentration, *f*_crit_ — may vary spatially (see below), which could help account for *in vivo* patterns of retinal degeneration.

Rates of rod degeneration in human RP have not been studied in great detail. Thus far, histopathological examination of human RP retinas has revealed that rod degeneration is most severe in the mid-peripheral retina, with relative sparing of rods in the macula and far-periphery until later in the disease (Milam et al., 1998). It may be that this pattern varies depending upon the mutation involved and between individuals (*cf*. Huang et al., 2012, for which different spatial patterns of rod function loss occur in patients, all of whom have a mutation in the RPGR gene). The rate of decay of rod photoreceptors has also been shown to vary with retinal eccentricity in mouse and pig models of RP (Carter-Dawson et al., 1978; Li et al., 1998). Further, under healthy conditions, the RdCVF concentration at the centre of the retina (near *θ* = 0) is much lower (*f* (*θ, t*) *∼ O*(10^−5^)) than in the remainder of the retina (where *f* (*θ, t*) *∼ O*(0.1) to *O*(1), see Fig. 3A). Therefore, it is reasonable to assume that central retinal cones are able to cope with lower RdCVF concentrations than those toward the periphery, and hence that *f*_crit_ is also heterogeneous. To determine whether these heterogeneities could account for cone degeneration patterns in human RP, we formulate and solve something known as an *inverse problem*.

In an inverse problem we seek to determine the model input required to attain a known/desired output. In this case the known output is the target cone degeneration profile, *t*_degen_(*θ*), while the input is either the rate of mutation-induced rod loss profile, *ϕ*_*r*_(*θ*), or the TF threshold concentration profile, *f*_crit_(*θ*), with corresponding inverses denoted as 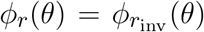 and 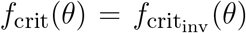 respectively. When searching for 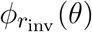, we hold the TF threshold concentration constant at *f*_crit_(*θ*) = *f*_crit_ = 3 × 10^−5^, while, when searching for 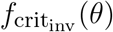, we hold the rate of mutation-induced rod loss constant at *ϕ*_*r*_(*θ*) = *ϕ*_*r*_ = 7.33 × 10^−2^. The constant value of *f*_crit_ is chosen to lie just below the minimum steady-state value of *f* (*θ*), such that, in the absence of rod loss, cones remain healthy, while the constant value of *ϕ*_*r*_ is chosen to be one hundred times higher than the value that can be inferred from measurements in the healthy human retina (Curcio et al., 1993), placing the timescale of the resultant cone loss on the order of decades, in agreement with *in vivo* RP progression rates (Hamel, 2006; Hartong et al., 2006).

We consider a range of target cone degeneration profiles, summarised in Table 3 and Fig. 5, which qualitatively replicate visual field loss Patterns 1A, 1B and 3 seen *in vivo* (and hence the corresponding *in vivo* cone degeneration patterns; taking the degeneration of the far-peripheral retina to occur in a radially symmetric manner in Pattern 3 — see Fig. 2 and Grover et al., 1998). We do not consider patterns of type 2 (to be explored in a future study) as these cannot be replicated by a 1D model (since the radial symmetry, assumed by the 1D model, is broken by variation in the azimuthal/circumferential direction). For each degeneration pattern we consider a set of sub-patterns to examine how this affects the shape of the inverses, allowing us to confirm that a modest change in the degeneration pattern results in a modest change in the inverses, while exploring both linear/piecewise linear profiles and more biologically realistic nonlinear (quadratic/cubic/exponential) patterns. We also consider a uniform target cone degeneration profile for comparison.

**Table 3.**
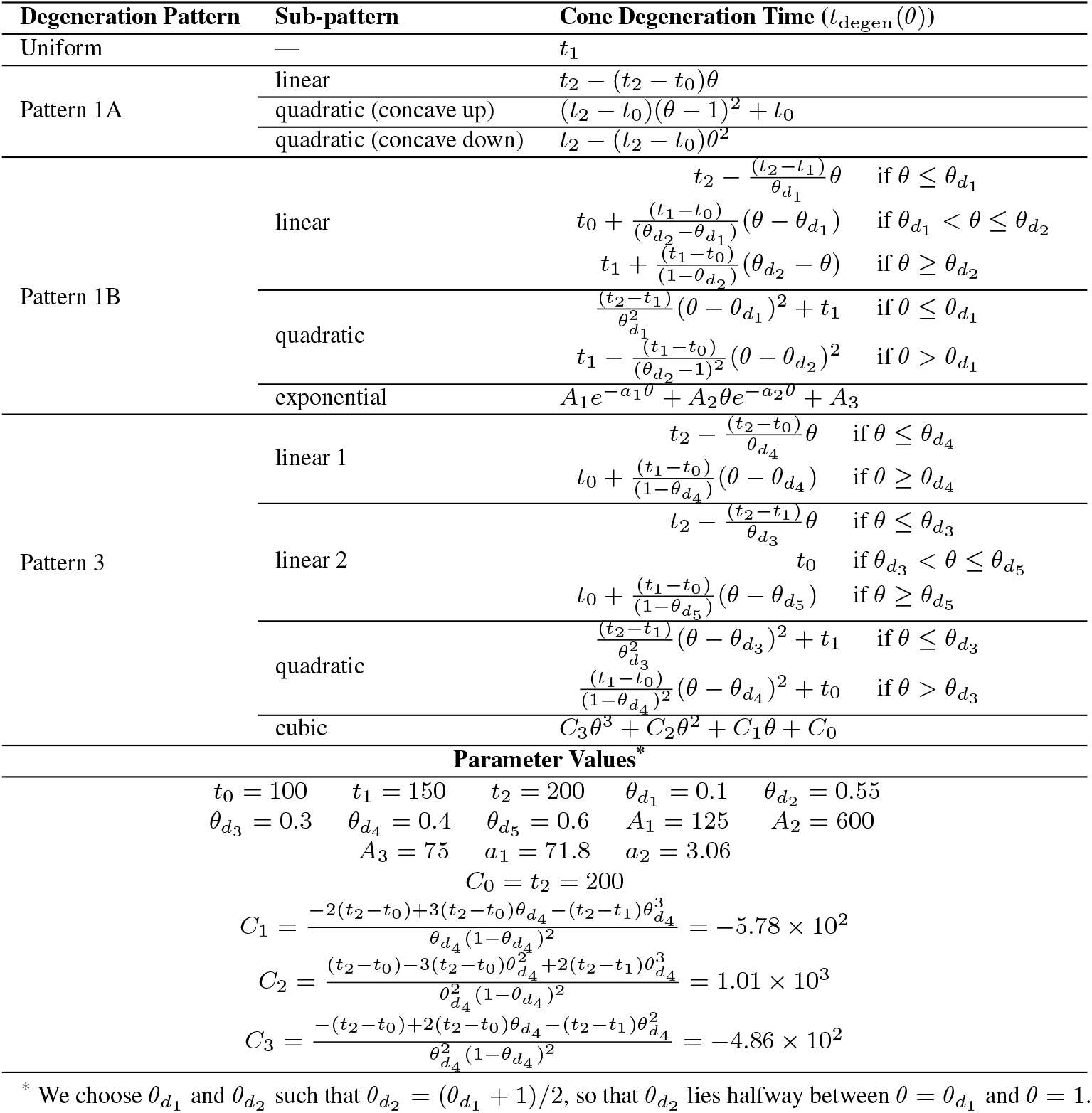
Target cone degeneration profiles, *t*_degen_(*θ*).

For each pattern, we consider the effect of two (biologically realistic) scalings for the rate of TF production by rods, *α*, and the rate of TF consumption by cones, *β*, upon the inverse profiles: (i) Scaling 1 — for which *α* = 7.01 × 10^4^ and *β* = 1.79 × 10^6^ as in Roberts (2022); and (ii) Scaling 2 — for which *α* = 7.01 × 10^2^ and *β* = 1.79 × 10^4^. Under Scaling 1, production and consumption of TF dominate over decay (with rate constant *η*), such that decay has a negligible effect upon the TF profile and model behaviour. Under Scaling 2, TF production and consumption occur at a similar rate to decay, such that they balance each other, resulting in a different TF profile and model behaviour (see Fig. 3A and C). As discussed in Roberts (2022), none of *α, β* or *η* have been measured. The decay rate, *η*, was chosen to match the measured decay rate of proteins in living human cells (Eden et al., 2011). Under Scaling 1, the consumption rate, *β*, is chosen such that it dominates over the decay rate (being a factor *ϵ*^−1^ = *O*(10^2^) larger), while the production rate, *α*, is chosen to balance consumption (see the Analytical Inverse section). This is a sensible scaling as it is likely that cones consume RdCVF at a much faster rate than that at which it decays. It is possible, however, that cones consume RdCVF at a similar rate to its decay rate, which is the scenario we consider in Scaling 2; reducing *α* and *β* by a factor of 100 (*∼ ϵ*^−1^) to bring consumption and production into balance with decay (see the Analytical Inverse section).

We solve the inverse problem both analytically and numerically (computationally), as described in the Analytical Inverse and Numerical Inverse sections below. Analytical approximations are computationally inexpensive and provide deeper insight into model behaviour, while numerical solutions, though computationally intensive, are more accurate.

#### 2.3.1 Analytical Inverse

Less mathematically inclined readers may wish to skip over the following derivation and proceed to the resulting Eqs. (6)–(11) and surrounding explanatory text. To derive analytical (algebraic) approximations for the inverses, 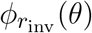 and 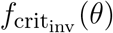, we perform an asymptotic analysis, seeking the leading order behaviour of Eqs. (1)–(5). In other words, we are simplifying our equations, making it possible to solve them algebraically (by hand), by only including those terms (corresponding to specific biological processes, e.g. TF production) which dominate the behaviour of the solution, where the method known as ‘asymptotic analysis’ enables us to rationally identify these dominant terms. Proceeding as in Roberts (2022) (where we considered a steady-state problem), we choose *ϵ* = *O*(10^−2^) and scale the parameters *η* = *ϵ*^−1^*η*^*′*^ and 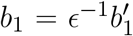, introducing the new scaling 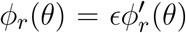, as we study the time-dependent problem here (where dashes ^*′*^ denote scaled variables and parameters). We consider two possible (biologically realistic) scalings on *α* and *β*: (i) Scaling 1 — for which *α* = *ϵ*^−2^*α*^*′*^ and *β* = *ϵ*^−3^*β*^*′*^ as in Roberts (2022) (corresponding to *α* = 7.01 × 10^4^ and *β* = 1.79 × 10^6^); and (ii) Scaling 2 — for which *α* = *ϵ*^−1^*α*^*′*^ and *β* = *ϵ*^−2^*β*^*′*^ (corresponding to *α* = 7.01 × 10^2^ and *β* = 1.79 × 10^4^). All remaining parameters are assumed to be *O*(1). We also scale the dependent variable 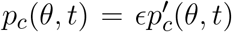, and assume *f* (*θ, t*) = *O*(1) and *p*_*r*_(*θ, t*) = *O*(1).

Applying the above scalings and dropping the dashes (working with the scaled versions of the variables and parameters, but omitting the dashes ^*′*^ for notational convenience), Eqn. (2) becomes

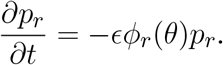

Thus, on this (fast) timescale, the rod density is constant. Since we are interested in the timescale upon which rods degenerate, we scale time as *t*^*′*^ = *ϵt* such that the decay term enters the dominant balance. Thus, on this slow timescale, after dropping the dashes, we have that

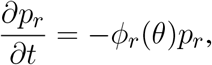

such that, at leading order, 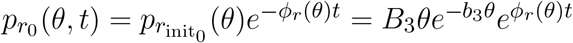.

We are interested here in the regime in which cones have not yet degenerated, thus we assume the leading order cone density remains constant at 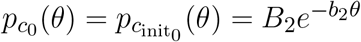.

Applying Scaling 1 and the slow timescale to Eqn. (1) we obtain

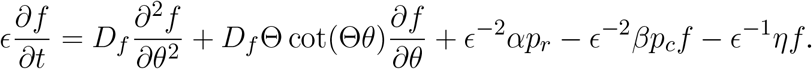

Since the TF dynamics occur on a faster timescale than mutation-induced rod loss, we make a quasi-steady-state approximation (QSSA), assuming that the TF concentration instantaneously takes its steady-state profile, for any given rod density profile, as the rods degenerate (*ϵ∂*_*t*_*f ∼* 0). Thus, at leading order, we obtain

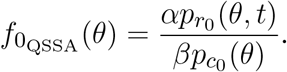

Rearranging this expression and assuming that cone degeneration initiates when 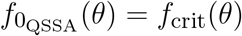, we obtain the cone degeneration time profile,

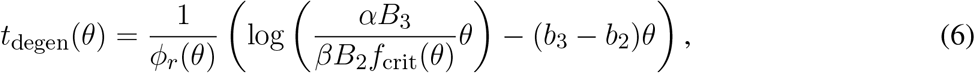

the inverse mutation-induced rod degeneration rate profile,

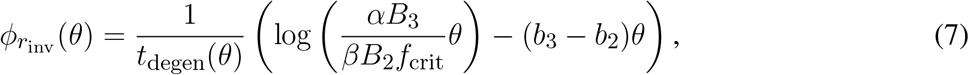

and the inverse TF threshold concentration profile,

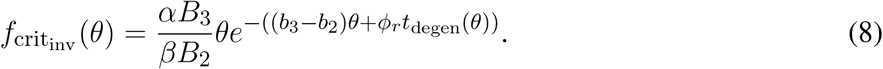

Alternatively, if we apply Scaling 2 and the slow timescale to Eqn. (1) we obtain

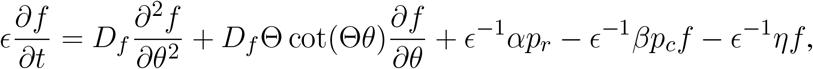

with the TF decay term, *ηf*, now entering the dominant balance. Applying the QSSA and proceeding as above we find

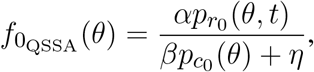

with cone degeneration time profile,

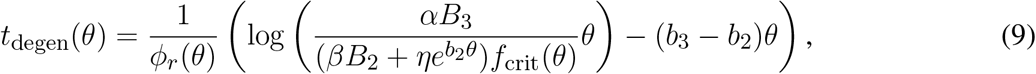

inverse mutation-induced rod degeneration rate profile,

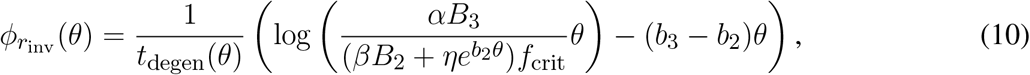

and inverse TF threshold concentration profile,

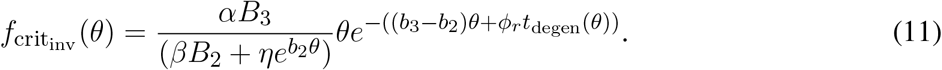

These equations reveal how the inverses, 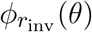 and 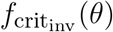, are influenced by our choices for fixed values of *f*_crit_ and *ϕ*_*r*_, respectively. As can be seen from Eqs. (7) and (10), 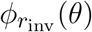 is inversely and monotonically related to *f*_crit_, such that as *f*_crit_ increases, 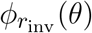 decreases. Similarly, 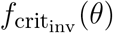 and *ϕ*_*r*_ are inversely and monotonically related in Eqs. (8) and (11), such that as *ϕ*_*r*_ increases, 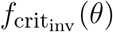 decreases. Lastly, as would be expected intuitively, *t*_degen_(*θ*), 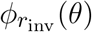 and 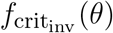 all increase monotonically with increasing TF production, *α*, and decrease monotonically with increasing TF consumption, *β*, and TF decay *η* (Eqs. (6)–(8) and (9)–(11)).

#### 2.3.2 Numerical Inverse

The numerical inverse is calculated by repeatedly solving the forward problem (Eqs. (1)–(5)) for different values of the input (*ϕ*_*r*_(*θ*) or *f*_crit_(*θ*)), with the aim of converging upon the inverse (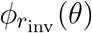 or 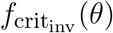). To find 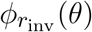 we use the Matlab routine fminsearch (which uses a simplex search method), while to obtain 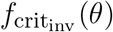 the Matlab routine patternsearch (which uses an adaptive mesh technique) was found to be more effective. In both cases the objective function (the quantity we are seeking to minimise) was taken as the sum of squares of the difference between the target cone degeneration profile, *t*_degen_(*θ*), and the contour described by 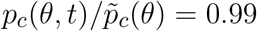 (along which cone degeneration is deemed to have initiated). Eqs. (1)–(5) were solved at each iteration as described in the Numerical Solutions section. Numerical inverses were calculated only at those locations (eccentricities) where the analytical inverse failed to generate a *t*_degen_(*θ*) profile matching the target profile, the analytical inverse being assumed to hold at all other eccentricities.

## 3 RESULTS

We begin by calculating the cone degeneration profiles, *t*_degen_(*θ*), in the case where both the rate of mutation induced rod degeneration, *ϕ*_*r*_, and the TF threshold concentration, *f*_crit_, are spatially uniform (or piecewise constant). We set the standard value for *ϕ*_*r*_ = 7.33 × 10^−2^ and consider the subcases (i) *f*_crit_ = 3 × 10^−5^ for 0 *≤ θ ≤* 1 (Fig. 4A), and (ii) *f*_crit_ = 0.3 for *θ >* 0.13 while *f*_crit_ = 3 × 10^−5^ for *θ ≤* 0.13 (Fig. 4B), as were explored in Roberts (2022). These subcases correspond to the situation in which the TF threshold concentration lies beneath the minimum healthy TF value at all retinal locations (i), and the situation in which foveal cones are afforded special protection compared to the rest of the retina, such that they can withstand lower TF concentrations (ii). For notational simplicity, we shall refer to subcase (ii) simply as *f*_crit_ = 0.3 in what follows. As with Figs. 6–9, we consider both Scaling 1 and Scaling 2 (see Inverse Problem) on the rate of TF production by rods, *α*, and the rate of TF consumption by cones, *β*, calculating both analytical and numerical solutions.

**Figure 4.**
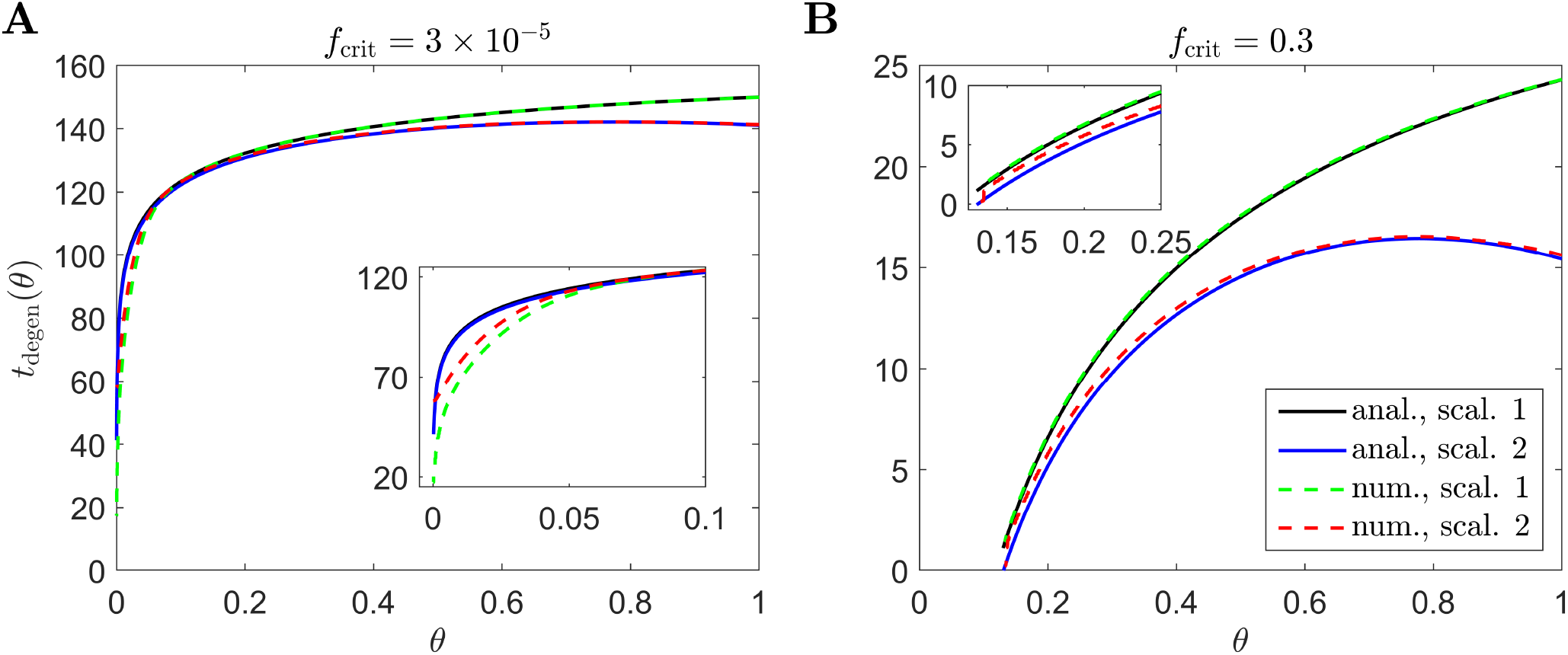
Cone degeneration profiles. Graphs show the time, *t*_degen_(*θ*), at which cones degenerate due to RdCVF deprivation, with constant rate of mutation-induced rod degeneration, *ϕ*_*r*_ = 7.33 × 10^−2^, and constant TF threshold concentrations: *f*_crit_ = 3 × 10^−5^ **(A)** and *f*_crit_ = 0.3 **(B)**. The solid black and dashed green curves correspond to Scaling 1 (*α* = 7.01 × 10^4^ and *β* = 1.79 × 10^6^), while the solid blue and dashed red curves correspond to Scaling 2 (*α* = 7.01 × 10^2^ and *β* = 1.79 × 10^4^). The black and blue solid curves are analytical approximations, obtained by plotting Eqs. (6) and (9) respectively, while the green and red dashed curves are 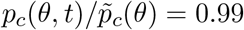 contours, obtained by solving Eqs. (1)–(5) using the method of lines with 401 mesh points. **(A)** simulation spans *∼* 17.7 years in dimensional variables; **(B)** simulation spans *∼* 2.8 years in dimensional variables. Insets show magnified portions of each graph. Cone degeneration initiates at the fovea (*θ* = 0) in **(A)** and at *θ* = 0.13 in **(B)**, spreading peripherally (rightwards) in both cases. Degeneration occurs earlier in **(B)** than in **(A)** and for Scaling 2 than for Scaling 1 (except near the fovea in **(A)**). Remaining parameter values as in Table 2.

Cone degeneration initiates at the fovea (*θ* = 0) in Fig. 4A and at *θ* = 0.13 in Fig. 4B, spreading peripherally (rightwards) in both cases, while degeneration also initiates at the ora serrata (*θ* = 1) under Scaling 2 in both Fig. 4A and Fig. 4B, spreading centrally. Degeneration occurs earlier in Fig. 4B than in Fig. 4A and earlier for Scaling 2 than for Scaling 1 (except near the fovea in Fig. 4A). Numerical and analytical solutions agree well, only diverging close to the fovea in Fig. 4A, where the analytical solution breaks down. None of these patterns of degeneration match those seen *in vivo* (see Fig. 2).

In Figs. 6–9 we calculate the 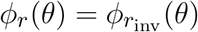 and 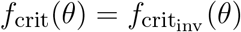 profiles required to qualitatively replicate the cone degeneration profiles, *t*_degen_(*θ*), observed *in vivo* (Fig. 5), by solving the associated inverse problems (see Inverse Problem). As noted in the Inverse Problem section, when searching for 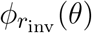, we hold the TF threshold concentration constant at *f*_crit_(*θ*) = *f*_crit_ = 3 × 10^−5^, while, when searching for 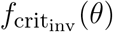, we hold the rate of mutation-induced rod loss constant at *ϕ*_*r*_(*θ*) = *ϕ*_*r*_ = 7.33 × 10^−2^. Analytical inverses are plotted across the domain (0 *≤ θ ≤* 1), while numerical inverses are calculated and plotted only at those locations (eccentricities) where the analytical inverse fails to generate a *t*_degen_(*θ*) profile matching the target profile (as determined by visual inspection, the *t*_degen_(*θ*) and target profiles being visually indistinguishable outside of these regions).

**Figure 5.**
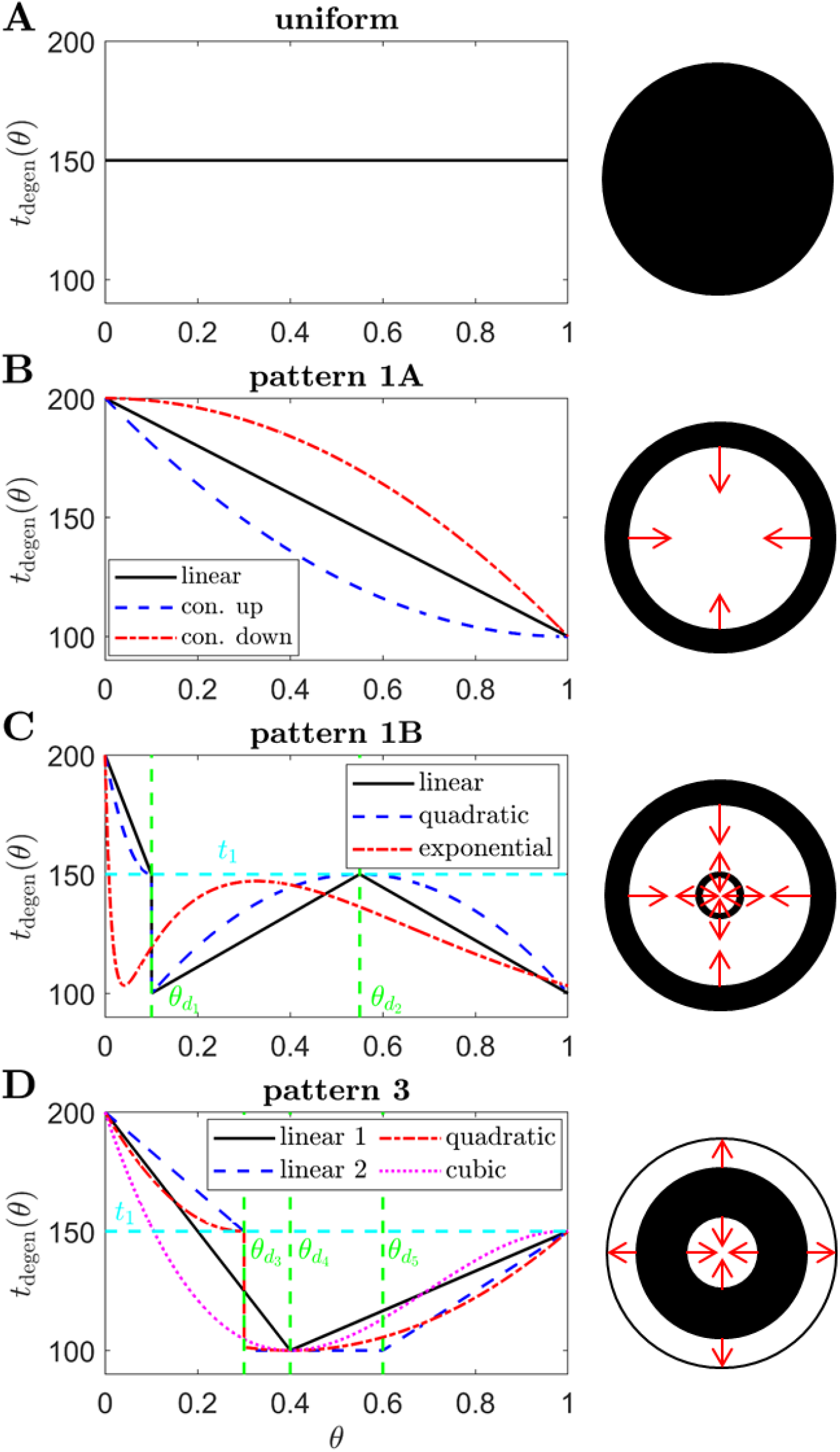
Target cone degeneration profiles. Panels (left) show cone degeneration profiles, *t*_degen_(*θ*), qualitatively replicating typical spatio-temporal patterns of visual field loss in RP: **(A)** Uniform, **(B)** Pattern 1A, **(C)** Pattern 1B and **(D)** Pattern 3. Visual field loss patterns directly correspond to cone degeneration patterns in these radially symmetric cases. We seek to replicate these patterns by finding appropriate 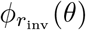 and 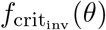 profiles in Figs. 6–9. Diagrams on the right show the corresponding 2D patterns of visual field loss — white regions: preserved vision, black regions: scotomas (blind spots), and red arrows: direction of scotoma propagation. Parameters: *t*_0_ = 100 (*∼* 11.0 years), *t*_1_ = 150 (*∼* 16.6 years), *t*_2_ = 200 (22.1 years, 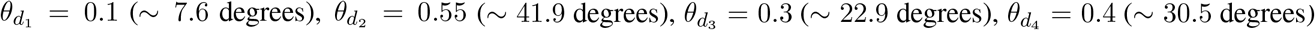 and 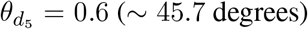. Cone degeneration profile formulas and parameters are given in Table 3. Remaining parameter values as in Table 2.

In Fig. 6 we calculate inverses for a Uniform degeneration profile. While this pattern is not typically observed in humans, we consider this case as a point of comparison with the non-uniform patterns explored in Figs. 7–9. Both inverses, 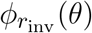 and 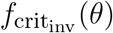, are monotone increasing for Scaling 1, and increase initially for Scaling 2 before reaching a maximum and decreasing toward the ora serrata (at *θ* = 1). Consequently, Scaling 1 and 2 inverses, while close near the fovea (*θ* = 0), diverge toward the ora serrata, this effect being more prominent for 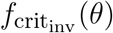. The inverse profiles have a similar shape to the *t*_degen_(*θ*) profiles in Fig. 4 (see Discussion). Numerical solutions reveal lower values of the inverses near the fovea, where the analytical approximations break down.

**Figure 6.**
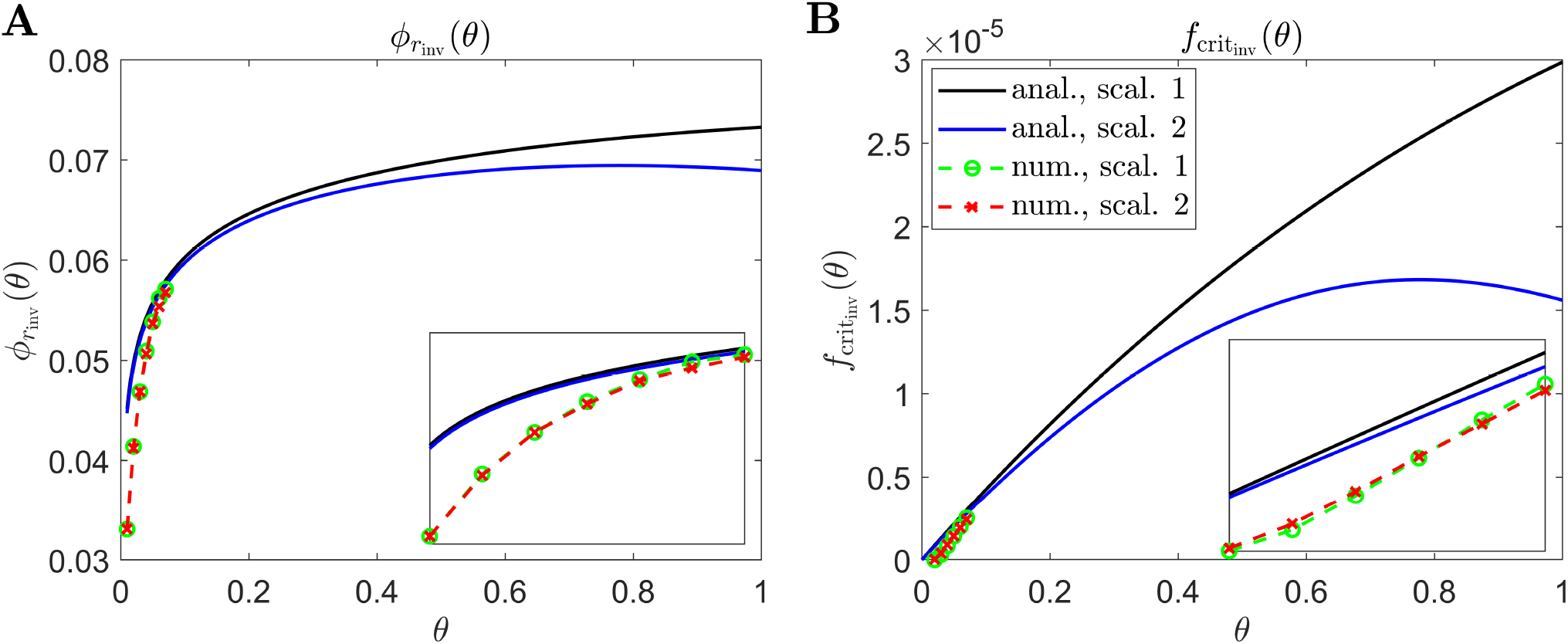
Inverse mutation-induced rod degeneration rate and TF threshold concentration — Uniform target cone degeneration profile. **(A)** inverse mutation-induced rod degeneration rate, 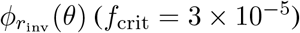; **(B)** inverse TF threshold concentration, 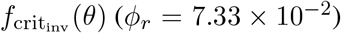. The solid black and dashed green curves correspond to Scaling 1 (*α* = 7.01 × 10^4^ and *β* = 1.79 × 10^6^), while the solid blue and dashed red curves correspond to Scaling 2 (*α* = 7.01 × 10^2^ and *β* = 1.79 × 10^4^). The black and blue solid curves are analytical approximations to the inverses, obtained by plotting Eqs. (7) and (10) respectively **(A)**, and Eqs. (8) and (11) respectively **(B)**. The green and red dashed curves are numerical inverses, obtained by using the Matlab routines fminsearch **(A)** and patternsearch **(B)** to calculate the *ϕ*_*r*_ and *f*_crit_ profiles for which the contour described by 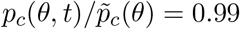 matches the target cone degeneration profile, *t*_degen_(*θ*). Eqs. (1)–(5) were solved at each iteration using the method of lines, with 101 mesh points. Insets show magnified portions of each graph. Numerical inverses are calculated and plotted only at those locations (eccentricities) where the analytical inverse fails to generate a *t*_degen_(*θ*) profile matching the target profile. Inverses are monotone increasing for Scaling 1, and increase initially for Scaling 2 before reaching a maximum and decreasing toward the ora serrata (*θ* = 1). Numerical solutions reveal lower values of the inverses near the fovea (*θ* = 0) than the analytical approximations suggest. Cone degeneration profile formulas and parameters are given in Table 3. Remaining parameter values as in Table 2.

**Figure 7.**
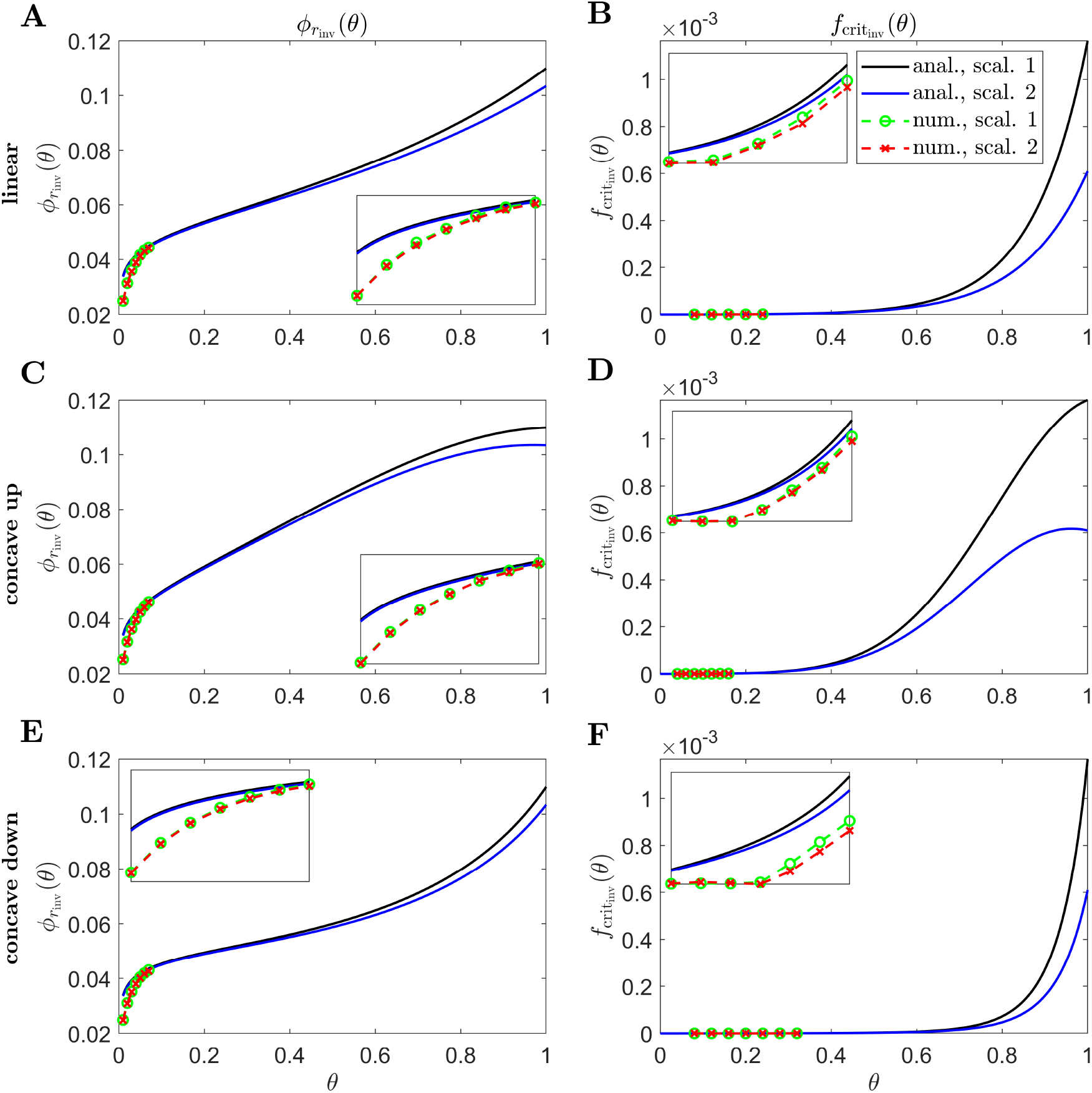
Inverse mutation-induced rod degeneration rate and TF threshold concentration — Pattern 1A target cone degeneration profiles. **(A), (C)** and **(E)** inverse mutation-induced rod degeneration rate, 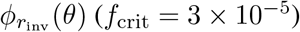; **(B), (D)** and **(F)** inverse TF threshold concentration, 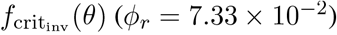. **(A)** and **(B)** linear target cone degeneration profile, *t*_degen_(*θ*); **(C)** and **(D)** concave up quadratic *t*_degen_(*θ*) profile; **(E)** and **(F)** concave down quadratic *t*_degen_(*θ*) profile. The solid black and dashed green curves correspond to Scaling 1 (*α* = 7.01 × 10^4^ and *β* = 1.79 × 10^6^), while the solid blue and dashed red curves correspond to Scaling 2 (*α* = 7.01 × 10^2^ and *β* = 1.79 × 10^4^). The black and blue solid curves are analytical approximations to the inverses, obtained by plotting Eqs. (7) and (10) respectively **(A), (C)** and **(E)**, and Eqs. (8) and (11) respectively **(B), (D)** and **(F)**. The green and red dashed curves are numerical inverses, obtained by using the Matlab routines fminsearch **(A), (C)** and **(E)**, and patternsearch **(B), (D)** and **(F)** to calculate the *ϕ*_*r*_ and *f*_crit_ profiles for which the contour described by 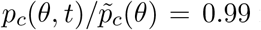 matches the target cone degeneration profile, *t*_degen_(*θ*). Eqs. (1)–(5) were solved at each iteration using the method of lines, with 26, 51 or 101 mesh points. Insets show magnified portions of each graph. Numerical inverses are calculated and plotted only at those locations (eccentricities) where the analytical inverse fails to generate a *t*_degen_(*θ*) profile matching the target profile. Inverses are monotone increasing functions for both scalings in **(A), (B), (E)** and **(F)**, and for Scaling 1 in **(C)** and **(D)**, while the inverses increase initially for Scaling 2 before reaching a maximum and decreasing toward the ora serrata (*θ* = 1) in **(C)** and **(D)**. Numerical solutions reveal lower values of the inverses near the fovea (*θ* = 0) than the analytical approximations suggest. Cone degeneration profile formulas and parameters are given in Table 3. Remaining parameter values as in Table 2.

Inverses for linear (Fig. 7A and B), concave up (quadratic) (Fig. 7C and D) and concave down (quadratic) (Fig. 7E and F) Pattern 1A degeneration profiles are shown in Fig. 7. Inverses are monotone increasing functions for both Scalings 1 and 2 in Fig. 7A,B,E and F, and for Scaling 1 in Fig. 7C and D, while the inverses increase initially for Scaling 2 before reaching a maximum and decreasing toward the ora serrata in Fig. 7C and D. Numerical solutions reveal lower values of the inverses near the fovea, where the analytical approximations break down.

Fig. 8 shows inverses for linear (Fig. 8A and B), quadratic (Fig. 8C and D) and exponential (Fig. 8E and F) Pattern 1B degeneration profiles. Inverses resemble vertically flipped versions of the *t*_degen_(*θ*) profiles in Fig. 5C (see Discussion). Numerical solutions reveal lower values of the inverses near the fovea, where the analytical approximations break down, and higher values in some regions away from the fovea in Fig. 8A–D. The discontinuities in the linear and quadratic cases are biologically unrealistic, though consistent with the idealised qualitative target cone degeneration patterns in Fig. 5C.

**Figure 8.**
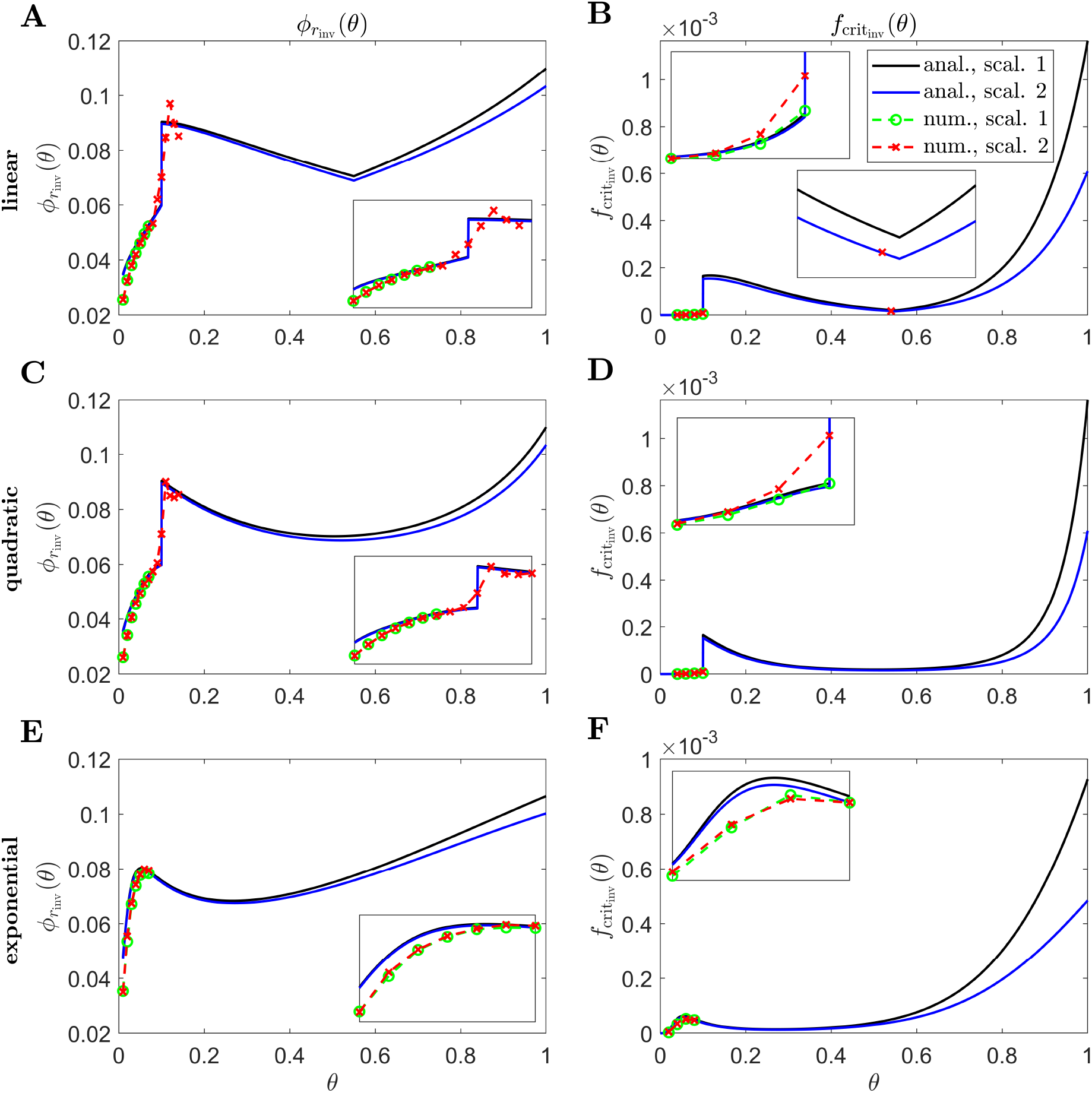
Inverse mutation-induced rod degeneration rate and TF threshold concentration — Pattern 1B target cone degeneration profiles. **(A), (C)** and **(E)** inverse mutation-induced rod degeneration rate, 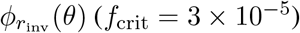; (**B**), **(D)** and **(F)** inverse TF threshold concentration, 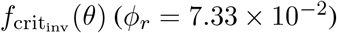. **(A)** and **(B)** linear target cone degeneration profile, *t*_degen_(*θ*); **(C)** and **(D)** quadratic *t*_degen_(*θ*) profile; **(E)** and **(F)** exponential *t*_degen_(*θ*) profile. The solid black and dashed green curves correspond to Scaling 1 (*α* = 7.01 × 10^4^ and *β* = 1.79 × 10^6^), while the solid blue and dashed red curves correspond to Scaling 2 (*α* = 7.01 × 10^2^ and *β* = 1.79 × 10^4^). The black and blue solid curves are analytical approximations to the inverses, obtained by plotting Eqs. (7) and (10) respectively **(A)**, (**C**) and **(E)**, and Eqs. (8) and (11) respectively **(B), (D)** and **(F)**. The green and red dashed curves are numerical inverses, obtained by using the Matlab routines fminsearch **(A), (C)** and **(E)**, and patternsearch **(B), (D)** and **(F)** to calculate the *ϕ*_*r*_ and *f*_crit_ profiles for which the contour described by 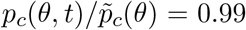 matches the target cone degeneration profile, *t*_degen_(*θ*). Eqs. (1)–(5) were solved at each iteration using the method of lines, with 51 or 101 mesh points. Insets show magnified portions of each graph. Numerical inverses are calculated and plotted only at those locations (eccentricities) where the analytical inverse fails to generate a *t*_degen_(*θ*) profile matching the target profile. Inverses resemble vertically flipped versions of the *t*_degen_(*θ*) profiles. Numerical solutions reveal lower values of the inverses near the fovea (*θ* = 0) than the analytical approximations suggest and higher values in some regions away from the fovea in **(A)**–**(D)**. Cone degeneration profile formulas and parameters are given in Table 3. Remaining parameter values as in Table 2.

In Fig. 9 we calculate inverses for linear 1 (Fig. 9A and B), linear 2 (Fig. 9C and D), quadratic (Fig. 9E and F) and cubic (Fig. 9G and H) Pattern 3 degeneration profiles. Inverses resemble vertically flipped versions of the *t*_degen_(*θ*) profiles in Fig. 5D (see Discussion). Numerical solutions reveal lower values of the inverses near the fovea, where the analytical approximations break down, and higher values in some regions away from the fovea in Fig. 9C–F and H. Similarly to Fig. 8, the discontinuities in the linear 2 and quadratic cases are biologically unrealistic, though consistent with the idealised qualitative target cone degeneration patterns in Fig. 5D.

**Figure 9.**
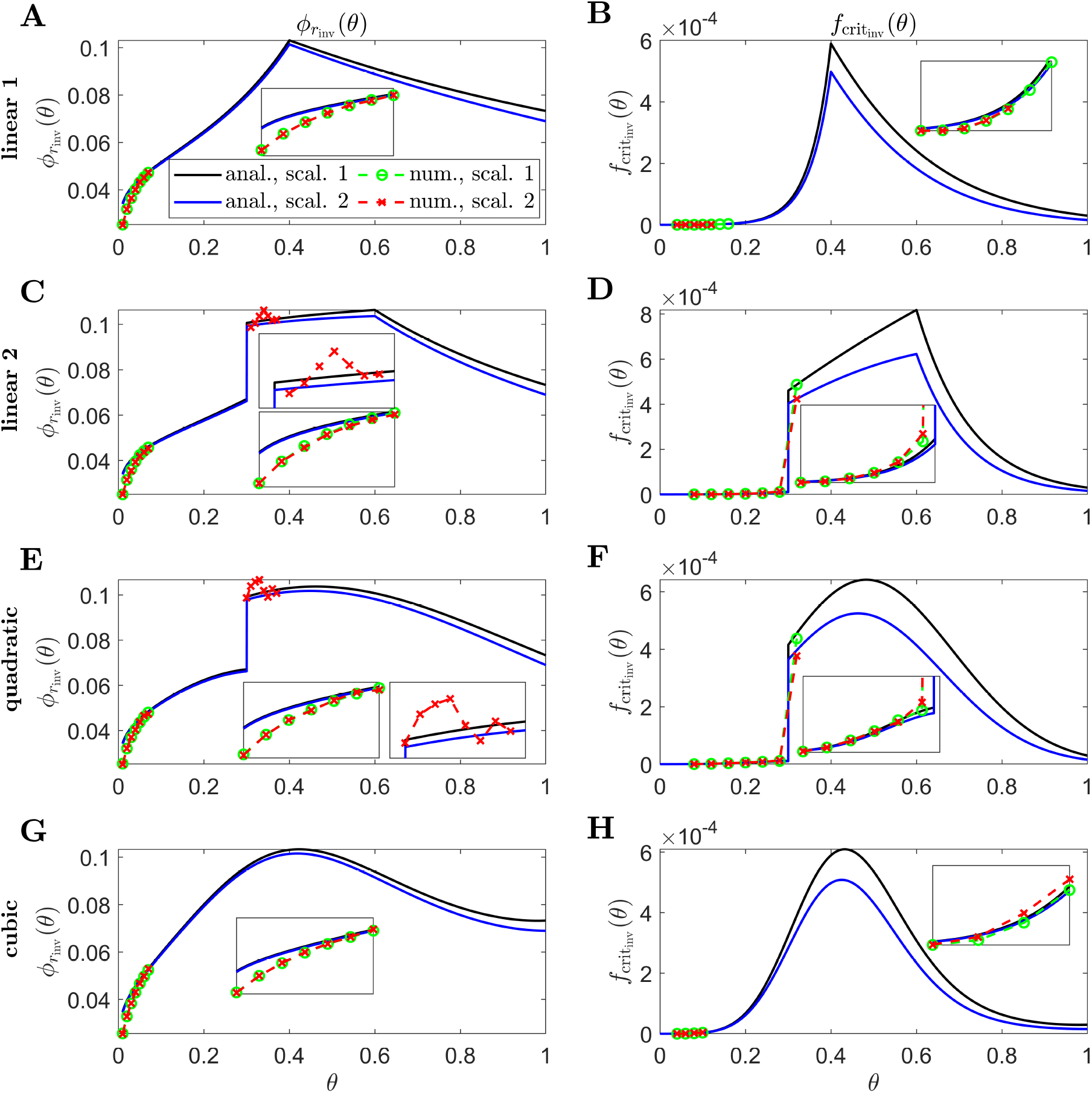
Inverse mutation-induced rod degeneration rate and TF threshold concentration — Pattern 3 target cone degeneration profiles. **(A), (C), (E)** and **(G)** inverse mutation-induced rod degeneration rate, 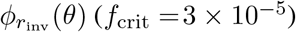; **(B), (D), (F)** and **(H)** inverse TF threshold concentration, 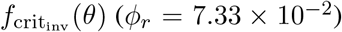. **(A)** and **(B)** linear 1 target cone degeneration profile, *t*_degen_(*θ*); **(C)** and **(D)** linear 2 *t*_degen_(*θ*) profile; **(E)** and **(F)** quadratic *t*_degen_(*θ*) profile; **(G)** and **(H)** cubic *t*_degen_(*θ*) profile. The solid black and dashed green curves correspond to Scaling 1 (*α* = 7.01 × 10^4^ and *β* = 1.79 × 10^6^), while the solid blue and dashed red curves correspond to Scaling 2 (*α* = 7.01 × 10^2^ and *β* = 1.79 × 10^4^). The black and blue solid curves are analytical approximations to the inverses, obtained by plotting Eqs. (7) and (10) respectively **(A), (C), (E)** and **(G)**, and Eqs. (8) and (11) respectively **(B), (D), (F)** and **(H)**. The green and red dashed curves are numerical inverses, obtained by using the Matlab routines fminsearch **(A), (C), (E)** and **(G)**, and patternsearch **(B), (D), (F)** and **(H)** to calculate the *ϕ*_*r*_ and *f*_crit_ profiles for which the contour described by 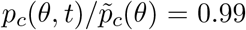 matches the target cone degeneration profile, *t*_degen_(*θ*). Eqs. (1)–(5) were solved at each iteration using the method of lines, with 26, 51 or 101 mesh points. Insets show magnified portions of each graph. Numerical inverses are calculated and plotted only at those locations (eccentricities) where the analytical inverse fails to generate a *t*_degen_(*θ*) profile matching the target profile. Inverses resemble vertically flipped versions of the *t*_degen_(*θ*) profiles. Numerical solutions reveal lower values of the inverses near the fovea (*θ* = 0) than the analytical approximations suggest and higher values in some regions away from the fovea in **(C)**–**(F)** and **(H)**. Cone degeneration profile formulas and parameters are given in Table 3. Remaining parameter values as in Table 2.

Lastly, in Fig. 10, we show simulation results of proportional cone loss for analytical and numerical 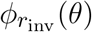 and 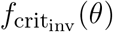, for Uniform (Scaling 1, Fig. 10A–D), concave up Pattern 1A (Scaling 1, Fig. 10E–H), linear Pattern 1B (Scaling 2, Fig. 10I–L) and quadratic Pattern 3 (Scaling 2, Fig. 10M–P) target degeneration profiles. Cone degeneration profiles generally show good agreement with the target *t*_degen_(*θ*) profiles. There is some divergence from *t*_degen_(*θ*) for the analytical inverses near the fovea and at discontinuous or nonsmooth portions of *t*_degen_(*θ*); this is mostly corrected by the numerical inverses. This correction is not perfect near the centre of the fovea, where cones still degenerate earlier than the target profiles. This occurs because it is necessary to replace the Heaviside step function in *λ*_2_(*f*) (see Eqn. (3)) with a hyperbolic tanh function to satisfy the smoothness requirements for the numerical solver, with the result that the initiation of cone degeneration is sensitive to the low TF concentrations (*f* (*θ, t*) *<* 10^−4^) in that region.

**Figure 10.**
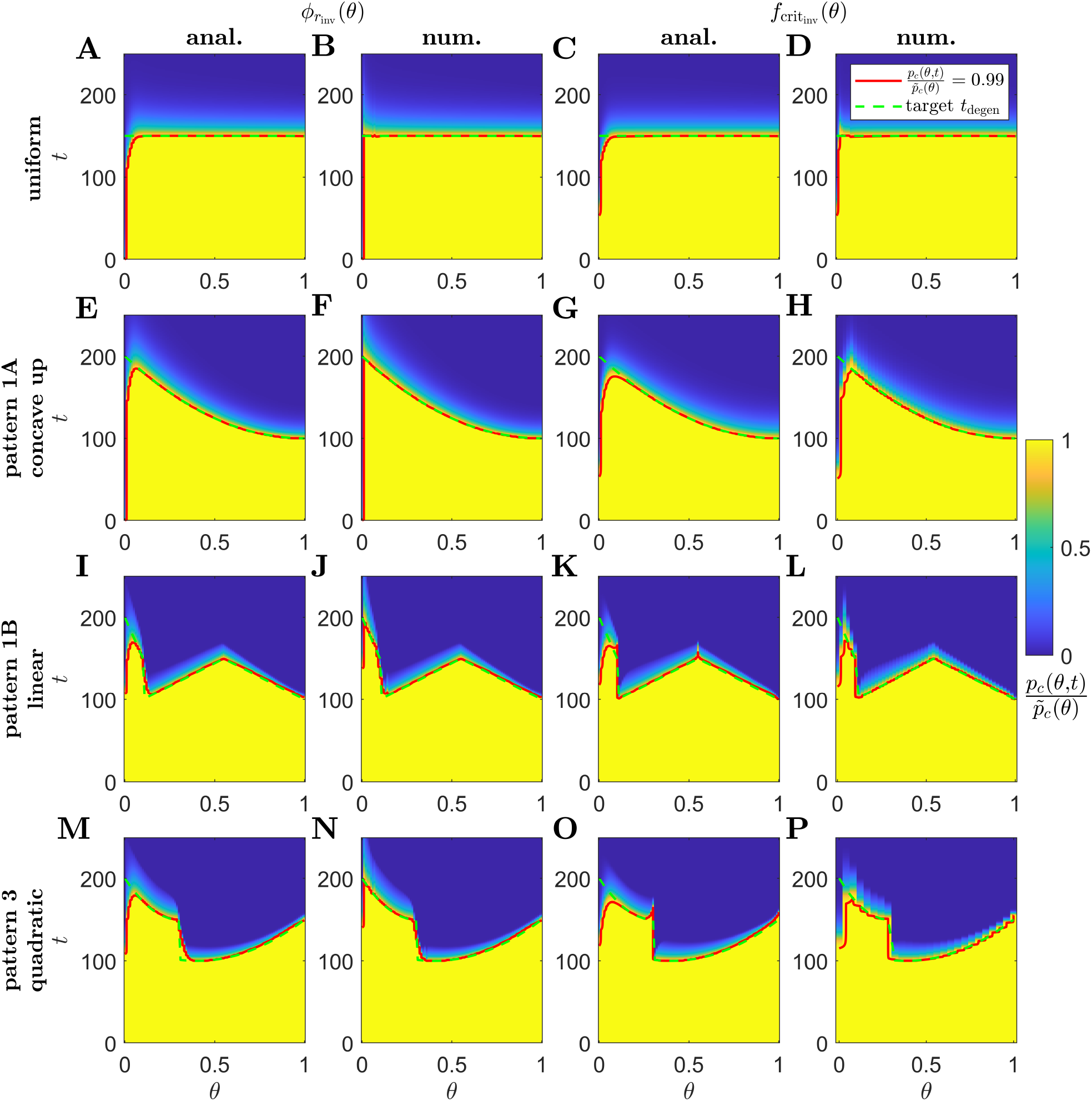
Simulations of proportional cone loss for a range of inverse mutation-induced rod degeneration rates and TF threshold concentrations. Plots show the proportion of cones remaining compared to local healthy values, 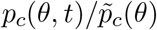, across space and over time. **(A), (E), (I)** and **(M)** analytical inverse mutation-induced rod degeneration rate, 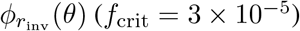; **(B), (F), (J)** and **(N)** numerical 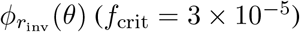; **(C), (G), (K)** and **(O)** analytical inverse TF threshold concentration, 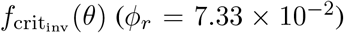; **(D), (H), (L)** and **(P)** numerical 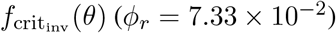. **(A)**–**(D)** Uniform target cone degeneration profile, *t*_degen_(*θ*), with Scaling 1 (*α* = 7.01 × 10^4^ and *β* = 1.79 × 10^6^); **(E)**–**(H)** Pattern 1A quadratic concave up *t*_degen_(*θ*) profile with Scaling 1; **(I)**–**(L)** Pattern 1B linear *t*_degen_(*θ*) profile with Scaling 2 (*α* = 7.01 × 10^2^ and *β* = 1.79 × 10^4^); **(M)**–**(P)** Pattern 3 quadratic *t*_degen_(*θ*) profile with Scaling 2. Eqs. (1)–(5) were solved using the method of lines, with 26, 51 or 101 mesh points. Analytical and numerical 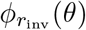 and 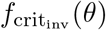 are as plotted in Figs. 6–9. Solid red curves denote the contours along which 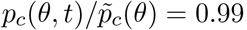, while dashed green curves show the target *t*_degen_(*θ*) profiles. Cone degeneration profiles generally show good agreement with the target *t*_degen_(*θ*) profiles. There is some divergence from *t*_degen_(*θ*) for the analytical inverses near the fovea (*θ* = 0) and at discontinuous or nonsmooth portions of *t*_degen_(*θ*); this is mostly corrected by the numerical inverses. Cone degeneration profile formulas and parameters are given in Table 3. Remaining parameter values as in Table 2.

## 4 DISCUSSION

The spatio-temporal patterns of retinal degeneration observed in human retinitis pigmentosa (RP) are well characterised; however, the mechanistic explanation for these patterns has yet to be conclusively determined. In this paper, we have formulated a one-dimensional (1D) reaction-diffusion partial differential equation (PDE) model (modified from Roberts, 2022) to predict RP progression under the trophic factor (TF) hypothesis. Using this model, we solved inverse problems to determine the rate of mutation-induced rod loss profiles, 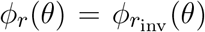, and TF threshold concentration profiles, 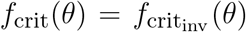, that would be required to generate spatio-temporal patterns of cone degeneration qualitatively resembling those observed *in vivo*, were the TF mechanism solely responsible for RP progression. In reality, multiple mechanisms (including oxidative damage and metabolic dysregulation, Punzo et al., 2009, 2012; Stone et al., 1999; Travis et al., 1991; Valter et al., 1998) likely operate in tandem to drive the initiation and propagation of retinal degeneration in RP. By using mathematics to isolate the TF mechanism, in a way that would be impossible to achieve experimentally, we are able to determine the conditions under which the TF mechanism alone would recapitulate known phenotypes. Having identified these conditions, this paves the way for future biomedical and experimental studies to test our predictions.

Other mechanisms may give rise to spatio-temporal patterns of retinal degeneration different from those predicted for the TF mechanism and may do so using fewer assumptions. For example, our previous work on oxygen toxicity in RP demonstrated that this mechanism can replicate visual field loss Pattern 1 (especially 1B) and the late far-peripheral degeneration stage of Pattern 3, without imposing heterogeneities on the rod decay rate or photoreceptor susceptibility to oxygen toxicity (Roberts et al., 2017, 2018). Further, we hypothesise that the toxic substance hypothesis (in which dying rods release a chemical which kills neighbouring photoreceptors) is best able to explain the early mid-peripheral loss of photoreceptors in Patterns 2 and 3, given the high density of rods in this region. In future work, we will explore the toxic substance and other hypotheses, ultimately combining them together in a more comprehensive modelling framework, aimed at explaining and predicting all patterns of retinal degeneration in RP.

Spatially uniform *ϕ*_*r*_(*θ*) and *f*_crit_(*θ*) profiles fail to replicate any of the *in vivo* patterns of degeneration (Fig. 4), showing that heterogenous profiles are required, all else being equal. Throughout this paper we have considered two scalings on the rate of TF production by rods, *α*, and the rate of TF consumption by cones, *β* (denoted as Scalings 1 and 2, see the Inverse Problem section for details). Under Scaling 1, the rod:cone ratio (Fig. 3B) dominates the model behaviour (see Eqn. (6)), leading to a monotone, central to peripheral pattern of degeneration, while under Scaling 2, the trophic factor decay term, *ηf*, enters the dominant balance (see Eqn. (9)), such that degeneration initiates at both the fovea and (later) at the ora serrata, the degenerative fronts meeting in the mid-/far-periphery (Fig. 4).

As discussed in the Inverse Problem section, the rate of mutation-induced rod loss, *ϕ*_*r*_(*θ*), is known to be spatially heterogeneous in humans with RP (Milam et al., 1998). The *ϕ*_*r*_(*θ*) profile predicted for Pattern 3 is consistent with the preferential loss of rods in the mid-peripheral retina noted by Milam et al. (1998) for human RP. A more extensive biomedical investigation is required to characterise quantitatively the diversity of *ϕ*_*r*_(*θ*) profiles across individuals and for different mutations. This would make it possible to determine if the *ϕ*_*r*_(*θ*) profiles predicted by our model for cone degeneration Patterns 1A and 1B are realised in human RP patients with those cone degeneration patterns. To the best of our knowledge, we are the first to suggest that the intrinsic susceptibility of cones to RdCVF deprivation, characterised in our models by the TF threshold concentration, *f*_crit_(*θ*), may vary across the retina. Assuming it does vary, what might account for this phenomenon? There is a precedent for special protection being provided to localised parts of the retina. For example, experiments in mice have found that production of basic fibroblast growth factor (bFGF) and glial fibrillary acidic protein (GFAP) is permanently upregulated along the retinal edges, at the ora serrata and optic disc, to protect against elevated stress in these regions (Mervin and Stone, 2002; Stone et al., 2005). Similarly, in the human retina, rods (though not cones) contain bFGF, with a concentration gradient increasing towards the periphery (Li et al., 1997, potentially explaining the relative sparing of rods often observed at the far-periphery). By analogy, we speculate that, in the human retina, cone protective factors may be upregulated at the fovea to compensate for the low RdCVF concentrations in that region, lowering the local value of *f*_crit_(*θ*). This hypothesis awaits experimental confirmation.

We solved the inverse functions, 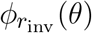 and 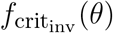, both analytically (algebraically) and numerically (computationally). Analytical solutions are approximations; however, they have the advantage of being easier to compute (increasing their utility for biomedical researchers) and provide a more intuitive understanding of model behaviour, while numerical solutions are more accurate, though computationally expensive. We calculated the inverses for a range of target cone degeneration profiles, consisting of a Uniform profile and profiles which qualitatively replicate those found *in vivo*: Pattern 1A, Pattern 1B and Pattern 3 (Pattern 2 being inaccessible to a 1D model; see Table 3 and Fig. 5).

The shapes of the inverse functions are determined partly by the rod and cone distributions, 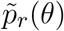 and 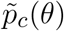, and partly by the target cone degeneration profile, *t*_degen_(*θ*) (see Eqs. (7), (8), (10) and (11)). As such, in the Uniform case (Fig. 6), the Scaling 1 inverse profiles take a similar shape to the rod:cone ratio (Fig. 3B), the inverses being lower towards the fovea to compensate for the smaller rod:cone ratio and hence lower supply of TF to each cone. The Scaling 2 inverse profiles follow a similar trend but decrease toward the ora serrata after peaking in the mid-/far-periphery due to the greater influence of the trophic factor decay term under this scaling. Interestingly, the shapes of these inverse profiles bear a striking resemblance to the cone degeneration profiles for spatially uniform *ϕ*_*r*_(*θ*) and *f*_crit_(*θ*) (Fig. 4). This is because lower values of the inverses are required to delay degeneration, in those regions where cones would otherwise degenerate earlier, to achieve a uniform degeneration profile. The inverse functions resemble vertically flipped versions of the target cone degeneration profiles for Patterns 1A, 1B and 3 (Figs. 7–9), this being more apparent for Patterns 1B and 3 due to their more distinctive shapes. This makes sense since lower inverse values are required for later degeneration times. Scaling 2 inverses typically lie below Scaling 1 inverses, compensating for the fact that degeneration generally occurs earlier under Scaling 2 than under Scaling 1 for any given *ϕ*_*r*_(*θ*) and *f*_crit_(*θ*).

Analytical inverses give rise to cone degeneration profiles that accurately match the target cone degeneration profiles, except near the fovea (centred at *θ* = 0, where the validity of the analytical approximation breaks down) and where the target *t*_degen_(*θ*) profile is nonsmooth or discontinuous (i.e. linear and quadratic Pattern 1B, and linear 1, linear 2 and quadratic Pattern 3; see Fig. 10 for examples). Numerical inverses improve accuracy in these regions, consistently taking lower values near the fovea, delaying degeneration where it occurs prematurely under the analytical approximation.

We have assumed throughout this study that at least one of *ϕ*_*r*_(*θ*) and *f*_crit_(*θ*) is spatially uniform. It is possible, however, that both vary spatially. In this case there are no unique inverses; however, if the profile for one of these functions could be measured experimentally, then the inverse problem for the remaining function could be solved as in this paper.

This work could be extended both experimentally and theoretically. Experimental and biomedical studies could measure how the rate of mutation-induced rod loss and TF threshold concentration vary with location in the retina, noting the spatio-temporal pattern of cone degeneration and comparing with the inverse 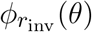 and 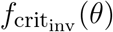 profiles predicted by our models. Curcio et al. (1993) have previously measured variation in the rate of rod loss in normal (non-RP) human retinas (where rods degenerated most rapidly in the central retina); a similar approach could be taken to quantify the rate of rod loss in human RP retinas. The parameter *f*_crit_ is less straightforward to measure. Léveillard et al. (2004) incubated cone-enriched primary cultures from chicken embryos with glutathione S-transferase-RdCVF (GST-RdCVF) fusion proteins, doubling the number of living cells per plate compared with GST alone. If experiments of this type could be repeated for a range of controlled RdCVF concentrations, then the value of *f*_crit_ could be identified. Determining the spatial variation of *f*_crit_(*θ*) in a foveated human-like retina may not be possible presently; however, the recent development of retinal organoids provides promising steps in this direction (Fathi et al., 2021; O’Hara-Wright and Gonzalez-Cordero, 2020). If organoids could be developed with a specialised macular region, mirroring that found *in vivo*, then the minimum RdCVF concentration required to maintain cones in health could theoretically be tested at a variety of locations. Further, the distribution of RdCVF, predicted in our models, could theoretically be measured in post-mortem human eyes using fluorescent immunohistochemistry, as was done for the protein neuroglobin by Ostojić et al. (2008) and Rajendram and Rao (2007), and perhaps also fluorescent immunocytochemistry as was done for bFGF by Li et al. (1997). In particular, it would be interesting to see if RdCVF concentration varies with retinal eccentricity as starkly as our model predicts, with extremely low levels in the fovea.

In future work, we will extend our mathematical model to two spatial dimensions, accounting for variation in the azimuthal/circumferential dimension (allowing us to capture radially asymmetric aspects of visual field loss Patterns 2 and 3, and to account for azimuthal variation in the rod and cone distributions), and use quantitative target cone degeneration patterns derived from SD-OCT imaging of RP patients (e.g. as in Escher et al., 2012). We will also adapt the model to consider animal retinas for which the photoreceptor distribution has been well characterised (e.g. rats, mice and pigs, Chandler et al., 1999; Gaillard et al., 2009; Ortín-Martínez et al., 2014).

In conclusion, we have formulated and solved a mathematical inverse problem to determine the rate of mutation-induced rod loss and TF threshold concentration profiles required to explain the spatio-temporal patterns of retinal degeneration observed in human RP. Inverse profiles were calculated for a set of qualitatively distinct degeneration patterns, achieving a close match with the target cone degeneration profiles. Predicted inverse profiles await future experimental verification.

## AUTHOR CONTRIBUTIONS

PAR: conceptualisation, methodology, software, validation, formal analysis, investigation, data curation, writing —- original draft, writing —- review and editing, visualisation, and project administration.

## ACKNOWLEDGMENTS

P.A.R. thanks Tom Baden for allowing the time to pursue this research. P.A.R. also thanks the reviewers for their helpful and insightful comments.

